# Structural and functional ramifications of antigenic drift in recent SARS-CoV-2 variants

**DOI:** 10.1101/2021.02.16.430500

**Authors:** Meng Yuan, Deli Huang, Chang-Chun D. Lee, Nicholas C. Wu, Abigail M. Jackson, Xueyong Zhu, Hejun Liu, Linghang Peng, Marit J. van Gils, Rogier W. Sanders, Dennis R. Burton, S. Momsen Reincke, Harald Prüss, Jakob Kreye, David Nemazee, Andrew B. Ward, Ian A. Wilson

## Abstract

The protective efficacy of neutralizing antibodies (nAbs) elicited during natural infection with SARS-CoV-2 and by vaccination based on its spike protein has been compromised with emergence of the recent SARS-CoV-2 variants. Residues E484 and K417 in the receptor-binding site (RBS) are both mutated in lineages first described in South Africa (B.1.351) and Brazil (B.1.1.28.1). The nAbs isolated from SARS-CoV-2 patients are preferentially encoded by certain heavy-chain germline genes and the two most frequently elicited antibody families (IGHV3-53/3-66 and IGHV1-2) can each bind the RBS in two different binding modes. However, their binding and neutralization are abrogated by either the E484K or K417N mutation, whereas nAbs to the cross-reactive CR3022 and S309 sites are largely unaffected. This structural and functional analysis illustrates why mutations at E484 and K417 adversely affect major classes of nAbs to SARS-CoV-2 with consequences for next-generation COVID-19 vaccines.

## INTRODUCTION

The COVID-19 pandemic has already lasted for over a year, but new infections are still escalating throughout the world. While several different COVID-19 vaccines have been deployed globally, a major concern is the emergence of antigenically distinct SARS-CoV- 2 variants. In particular, the B.1.351 (also known as 501Y.V2) lineage in South Africa (*1*) and B.1.1.28 lineage (and its descendant B.1.1.28.1, also known as P.1) in Brazil (*2*) have raised serious questions about the nature, extent and consequences of the antigenic drift observed in circulating SARS-CoV-2. The B.1.351 and B.1.1.28.1 lineages both share three mutations, namely K417N/T, E484K, and N501Y (which is also present in the UK B.1.1.7 lineage). A few B.1.1.7 genomes with the E484K mutation have also recently been detected (*3*). All of these mutations are located in the receptor binding site (RBS) of the receptor binding domain (RBD) of the spike (S) protein (Figure 1A). Two of these three mutations, K417N and E484K, decrease the neutralizing activity of sera as well as of monoclonal antibodies isolated from COVID-19 convalescent plasma and vaccinated individuals (*4–10*). Previous studies have shown that certain IGHV genes are highly enriched in the antibody response to SARS-CoV-2 infection, especially IGHV3-53 (*11–15*) and IGHV1-2 (*12, 16, 17*). IGHV3-53 and IGHV3-66, which differ by only one conservative substitution V12I, and IGHV1-2 are the most enriched IGHV genes used among 1,593 RBD antibodies from 32 studies (*12–43*) (Figure 1B). Thus, we have investigated the effects of these mutations on neutralization of SARS-CoV-2 by multi-donor class antibodies, and the consequences for current vaccines and therapeutics.

**Figure 1.**
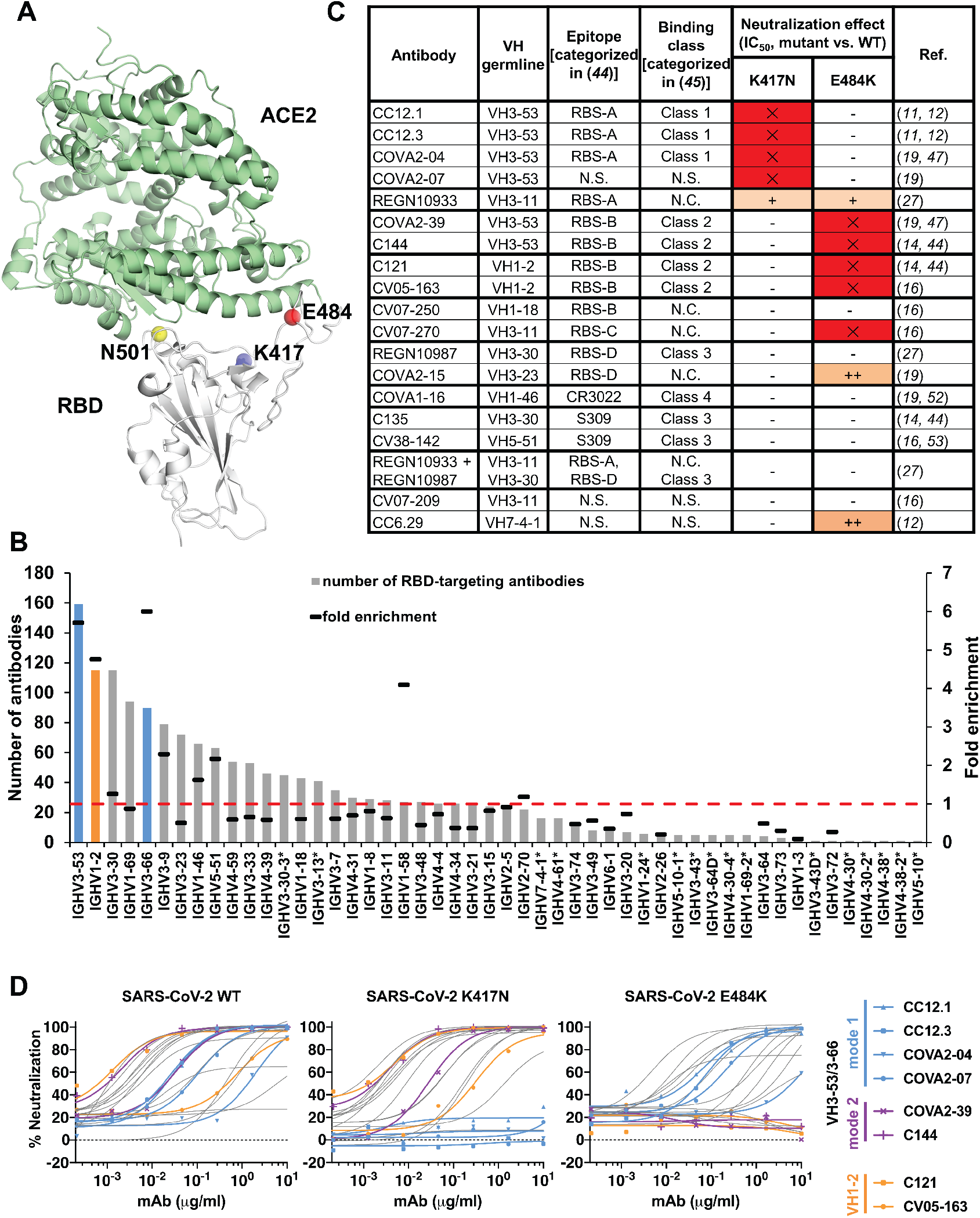
Emergent SARS-CoV-2 variants escape two major classes of neutralizing antibodies. **(A)** Emergent mutations (spheres) in the RBS of B.1.351 and B.1.1.28.1 lineages are mapped onto a structure of SARS-CoV-2 RBD (white) in complex with ACE2 (green) (PDB ID: 6M0J) (*61*). **(B)** Distribution of IGHV gene usage. The IGHV gene usage in 1,593 SARS-CoV-2 RBD-targeting antibodies (*12–43*) compared to healthy individuals (baseline) (*48*) is shown as bars. IGHV gene frequencies in healthy individuals that were not reported in (*48*) are shown with asterisks (*). Fold-enrichment of germlines used in SARS-CoV-2 antibodies over baseline is shown as black lines. A fold enrichment of one (red dashed line) represents no difference over baseline. The frequently used IGHV3-53 and IGHV3-66 genes are highlighted in blue, and IGHV1-2 in orange. Numbers of RBD- targeting antibodies encoded by each IGHV gene is shown as black lines. **(C)** Effect of single mutations on the neutralization activity of each neutralizing antibody. IC_50_ increases that are less than 10-fold are represented by “–”, between 10- and 100-fold as “+”, and greater than 100-fold as “++”. Results in red with “X” indicate no neutralization activity was detected at 10 μg/ml of IgG. N.C.: not categorized in the original studies. N.S.: No structure available. **(D)** Neutralization of pseudotyped SARS-CoV-2 virus and variants carrying K417N or E484K mutations. A panel of 18 neutralizing antibodies were tested, including four mode-1 IGHV3-53 antibodies (blue), two mode-2 IGHV3-53 antibodies (purple), and two IGHV1-2 antibodies (orange).

We tested the activity of a panel of 18 neutralizing antibodies isolated from COVID-19 patients or humanized mice against wild-type (Wuhan) SARS-CoV-2 pseudovirus, as well as single mutants K417N and E484K (Figure 1C). Neutralization of four and five antibodies out of the 18 tested antibodies were abolished by K417N and E484K, respectively. Strikingly, neutralization by all six highly potent IGHV3-53/3-66 antibodies that we tested was diminished for either K417N (binding mode 1) or E484K (binding mode 2) mutations (Figure 1C-D). In addition, neutralization by IGHV1-2 antibodies was strongly reduced in the E484K variant (Figure 1C-D). Consistently, binding of IGHV3-53/3-66 and IGHV1-2 antibodies to RBD was abolished by either K417N or E484K mutations (Figure S1).

We next examined all 49 SARS-CoV-2 RBD-targeting antibodies isolated from human patients with available structures. The epitopes of these antibodies on the RBD can be classified into six sites: RBS with four subsites RBS-A, B, C, and D; CR3022 site; and S309 site (Figure S2A). These epitopes assignments are related to the four classes in (*44*) (Figure 1C). Sixteen of 18 IGHV3-53/3-66 antibodies target RBS-A, which constitute the majority of RBS-A antibodies with reported structures (Figure 2A). All IGHV1-2 antibodies with available structures bind to the RBS-B epitope. A large fraction of the antibodies in these two main families interact with K417, E484 or N501 (Figure 2A). Almost all RBS-A antibodies interact extensively with K417 (and N501), whereas E484 is involved in interactions with most RBS-B and RBS-C antibodies. We also examined the buried surface area (BSA) of K417, E484, and N501 in the interface of these RBD-targeting antibodies (Figure S2B). The BSA confirmed why mutations at 417 and 484 affect binding and neutralization. Although several antibodies interact with N501, especially those targeting RBS-A, the N501 BSA is 30 Å^2^ or less, which is much smaller that the corresponding interactions with 417 and 484 (Figure S2B) (*45*). Antibodies targeting the RBS-D, S309 and CR3022 sites are minimally or not involved in interactions with any of these three mutated residues (Figure 2A-B).

**Figure 2.**
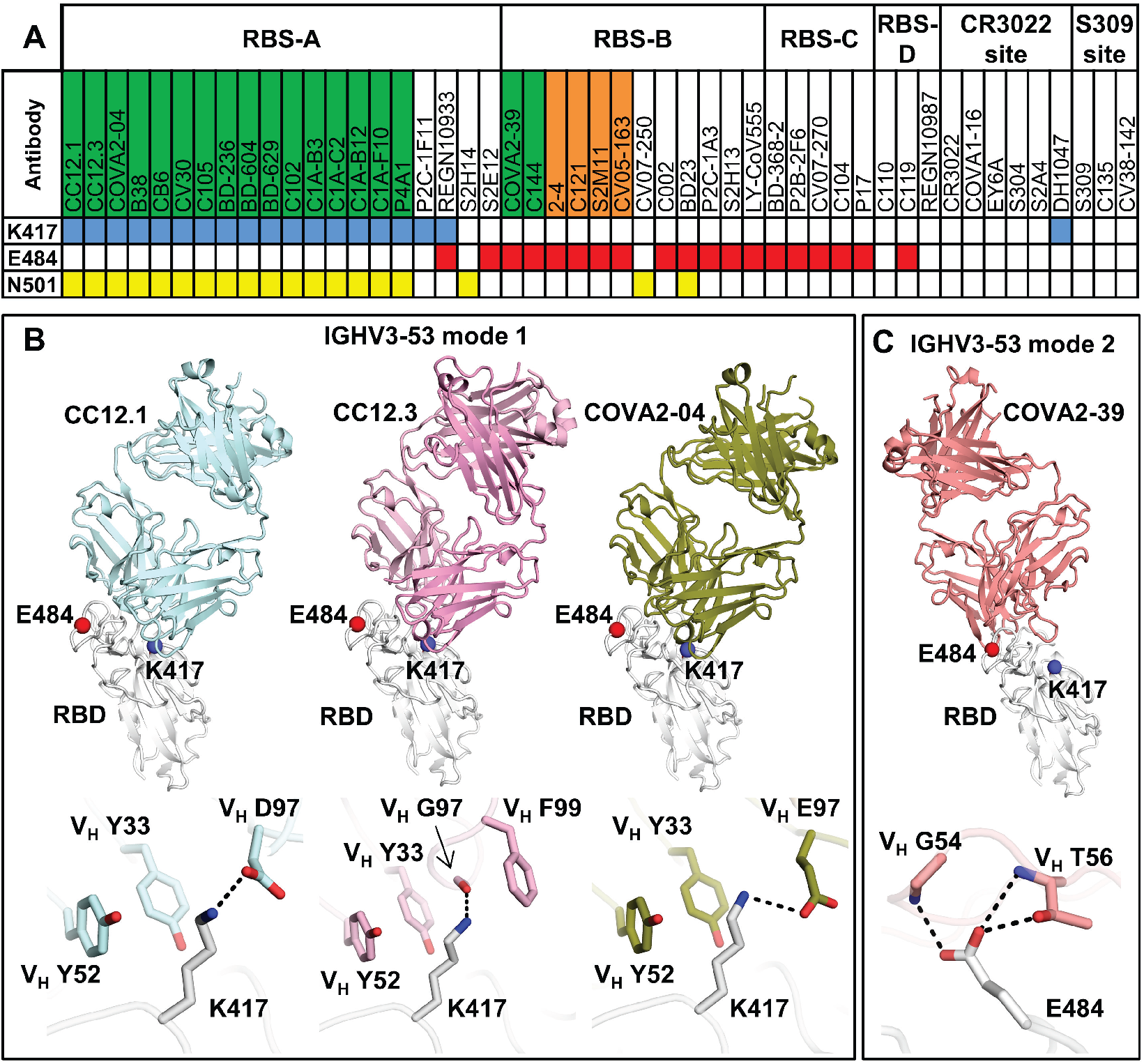
Antibody binding and structures to the wild-type SARS-CoV-2 RBS. **(A)** Antibodies making contact with RBD residues K417, E484 and N501 are represented by blue, red and yellow boxes, respectively (cutoff distance = 4 Å). Antibodies encoded by the most frequently elicited IGHV3-53/3-66 and IGHV1-2 in convalescent patients are shown in green and orange boxes, respectively. Antibodies are ordered by epitopes originally classified in (*46*) with an additional epitope RBS-D. Details of the epitope classifications are shown in Figure S2A. All available structures of RBD-targeting antibodies that were isolated from patients are analyzed. **(B-C)** Residues that are mutated in recently circulating variants are integral to the binding sites of IGHV3-53 antibodies. Representative structures are shown for **(B)** IGHV3-53 binding mode 1 [CC12.1 (PDB ID: 6XC3), CC12.3 (PDB ID: 6XC4) (*11*), and COVA2-04 (PDB ID: 7JMO) (*47*)] and **(C)** binding mode 2 [COVA2-39 (PDB ID: 7JMP) (*47*)]. The SARS-CoV-2 RBD is in white and Fabs in different colors. Residues K417 and E484 are represented by blue and red spheres, respectively. Hydrogen bonds and salt bridges are represented by black dashed lines.

Importantly, all of the IGHV3-53/3-66 antibodies that bind to the RBS-A epitope interact with K417 (and N501Y), consistent with our neutralization results (Figure 1C-D). Previously, we and others demonstrated that IGHV3-53/3-66 RBD antibodies can adopt two different binding modes (*44, 46*), which we now refer to as binding modes 1 and 2, with distinct epitopes and angles of approach to the receptor binding site (RBS) (Figure 2B, Figure S3). All known IGHV3-53/3-66 RBD antibodies with binding mode 1 have a short CDR H3 of < 15 amino acids and bind to the RBS-A epitope (*11, 15, 31*), while those with binding mode 2 contain a longer CDR H3 (≥ 15 amino acids) and target RBS-B (*44, 46, 47*). These dual binding modes enhance the recognition potential of this antibody family for the SARS-CoV-2 RBD, although 16 of 18 IGHV3-53/3-66 RBD antibodies with structural information adopt binding mode 1 (Figure S3). K417 is an important epitope residue for all 16 antibodies with IGHV3-53/3-66 binding mode 1 (Figure 2B, Figure S3). IGHV3-53 germline residues V_H_ Y33 and Y52 make hydrophobic interactions with the aliphatic component of K417, and its ε-amino group interacts with CDR H3 through a salt bridge (D97 or E97), hydrogen bond, or cation-π interactions (F99) (Figure 2B). K417N/T would diminish such interactions and, therefore, affect antibody binding and neutralization. This observation provides a structural explanation for K417N escape in all tested IGHV3- 53/3-66 antibodies with binding mode 1 (Figures 1C-D and 2B, Figure S3). In contrast, IGHV3-53/3-66 antibodies with binding mode 2 do not interact with RBD-K417 (Figure S3), but with E484. Their CDR H2 hydrogen bonds (H-bond) with E484 through main-chain and sidechain-mediated H-bond interactions (Figure 2C). Consistently, binding and neutralization of IGHV3-53/3-66 antibodies with binding mode 2 (Figure S3) are abolished by E484K, but not K417N (Figure 1C-D, Figure S1).

Among the IGHV genes used in RBD antibodies, IGHV1-2 is also highly enriched over the baseline frequency in the antibody repertoire of healthy individuals (*48*), and is second only to IGHV3-53/3-66 (Figure 1B). We compared three available structures of IGHV1-2 antibodies, namely 2-4 (*26*), S2M11 (*29*), and C121 (*44*). Similar to IGHV3-53/3-66 RBD antibodies, these IGHV1-2 antibodies also target the RBS, but to RBS-B. Despite being encoded by different IGK(L)V genes, 2-4 (IGLV2-8), S2M11 (IGKV3-20), and C121 (IGLV2-23) share a nearly identical binding mode and epitope (Figure 3A). Structural analysis reveals that the V_H_ ^26^GTFTG(Y)Y^33^, ^50^W(I)N/S(P)XSXGTX^58^, ^73^TS(I)S/T^76^ motifs are important for RBD binding (Figure S4A-D). Although only a small area of the epitope is conferred by the light chains of 2-4, S2M11, and C121, V_L_ residues 32 and 91 (n.b. also residue 30 for some antibodies) play an important role in forming a hydrophobic pocket together with V_H_ residues for binding RBD-F486, which we consider another key binding residue in such classes of antibodies (*46*) (Figure S4E-I). A recent study also showed that three other IGHV1-2 antibodies, 2-43, 2-15, and H4 bind in a similar mode, further highlighting structural convergence of IGHV1-2 antibodies in targeting the same RBD epitope (*49*). Importantly, all IGHV1-2 antibodies to date form extensive interactions with E484 (Figure 3A, Figure S2B). In particular, germline-encoded V_H_ Y33 and V_H_ N52 (somatically mutated to S52 in C121) and S54 are involved in polar interactions with the side chain of RBD-E484, where these H-bonds would be altered by substitution with Lys (Figure 3A) and diminish binding and neutralization of IGHV1-2 antibodies against E484K (Figure 1C-D, Figure S1). Consistent with the neutralization results, E484 has a large BSA when complexed with these antibodies (Figure S2B)

**Figure 3.**
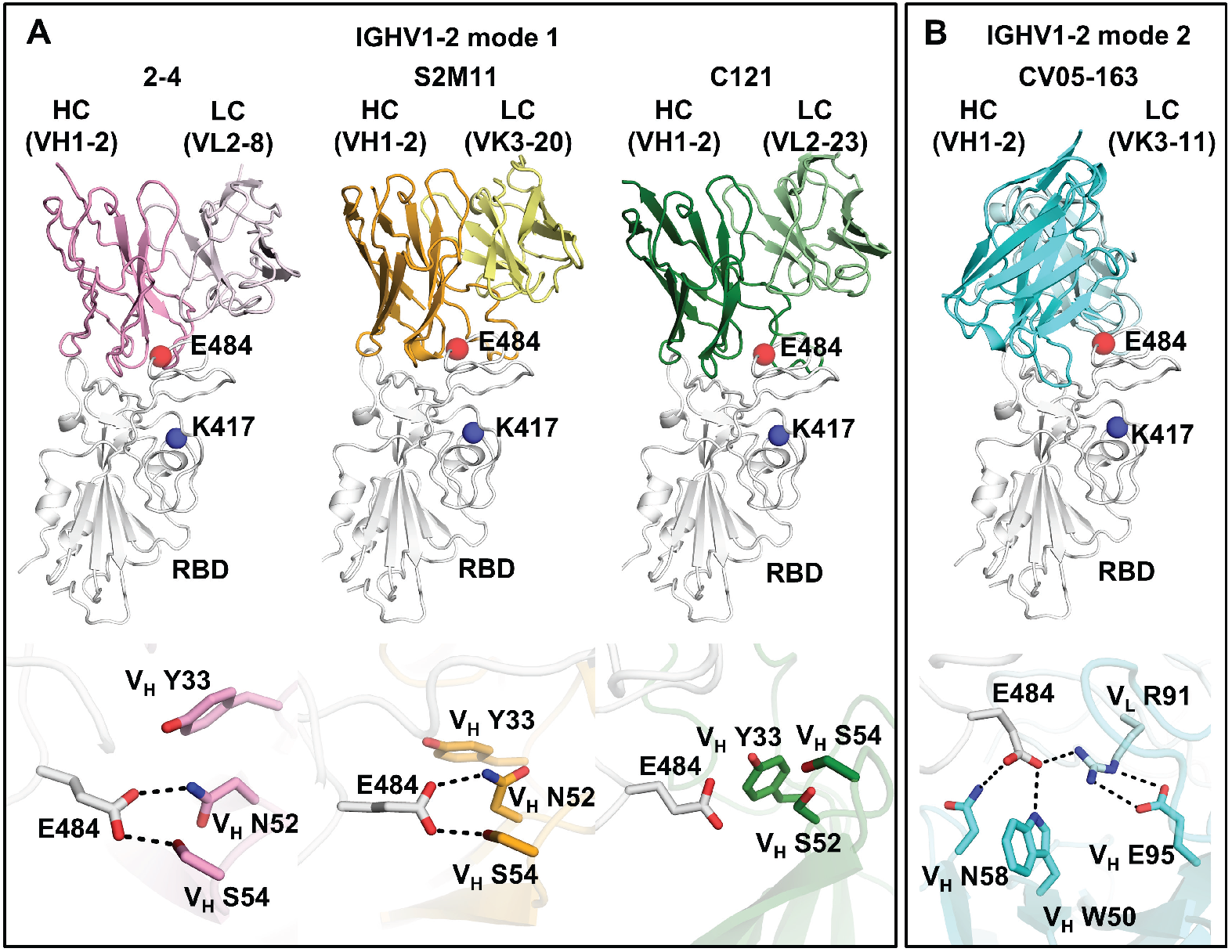
E484 is critical for RBD recognition of IGHV1-2 antibodies. Heavy and light chains of antibody 2-4 (PDB 6XEY) (*26*) are shown in pink and light pink, respectively, S2M11 (PDB 7K43) (*29*) in orange and yellow, and C121 (PDB 7K8X) (*44*) in dark and light green, and CV05-163 in cyan and light cyan. The RBD is shown in white. E484 and K417 are highlighted as red and blue spheres, respectively. Hydrogen bonds are represented by dashed lines. Hydrogen bonds are not shown in the panel of C121 due to the limited resolution (3.9 Å).

We previously isolated another IGHV1-2 antibody, CV05-163, from a COVID-19 patient (*16*). Interestingly, CV05-163 has no somatic mutations, and binds to SARS-CoV-2 RBD with a K_D_ of 0.2 nM as an IgG (*16*) and 45.3 nM as a Fab (Figure S5). CV05-163 also exhibited high neutralization potency with an IC_50_ of 16.3 ng/ml against authentic SARS-CoV-2 (*16*). Negative-stain electron microscopy (nsEM) of CV05-163 in complex with the SARS-CoV-2 S trimer illustrated that this antibody binds in various stoichiometries, including molar ratios of 1:1, 2:1, and 3:1 (Fab to S protein trimer), where RBDs in both up- and down-conformations can be accommodated by CV05-163 (Figure S6). We also determined a crystal structure of Fab CV05-163 in complex with SARS-CoV-2 RBD and Fab CR3022 to 2.25 Å resolution (Figure 3B, Figure S7, Tables S1 and S2) and show that it also binds to RBS-B. However, the binding orientation of Fab CV05-163 to the RBD (Figure 3B) is rotated 90° compared to other IGHV1-2 RBD antibodies 2-4, S2M11, and C121 (Figure 3A). CDR H2 forms three H-bonds as well as hydrophobic interactions with the RBD (Figure S7E). The non-templated nucleotide (N) additions in CDR H3 encode an ^100a^ALPPY^100e^ motif (Figures S7F and S8) that makes a major contribution to the RBD- interactions and promotes aromatic interactions between V_L_ Y32 and V_L_ Y49 and the RBD (Figure S9). All paratope residues on the light chain are encoded by IGKV3-11 (Figure S8), which form nine H-bonds and salt bridges as well as multiple hydrophobic interactions (Figure S7G-I). Of note, CV05-163 likely represents a public clonotype for IGHV1-2 RBD antibodies across patients (Figure S10). In contrast to the other IGHV1-2 antibodies with a canonical binding mode (2-4, S2M11, and C121) where four residues stack with RBD- F486, CV05-163 in this alternate binding mode interacts with RBD-F486 via only one residue (Figure S4E-G). Nevertheless, CV05-163 binds to a similar epitope as the other IGHV1-2 RBD antibodies (Figure 3). Importantly, CV05-163 also extensively interacts with RBD-E484 via H-bonds (V_H_ W50 and V_H_ N58) and a salt bridge (V_L_ R91) (Figure 3B) that explains why the binding and neutralization by CV05-163 were diminished by the E484K mutant (Figure 1C-D, Figure S1). As a result, IGHV1-2 antibodies [like IGHV3-53/66 (*47*)], can also engage the RBD in another example of two different binding modes, both of which are susceptible to escape by the E484K mutation, but not K417N (Figure 1C-D).

A further group of antibodies target the back side of the RBS on the opposite side of the RBS ridge (called RBS-C) (*46*). To date, structures of five antibodies isolated from COVID- 19 patients can be classified as binding to the RBS-C epitope: CV07-270 (*16*), BD-368-2 (*37*), P2B-2F6 (*17*), C104 (*44*), and P17 (*50*). All of these RBS-C antibodies also interact with E484 (Figure 4A), many with an Arg residue in CDR H3, suggesting that RBD-E484K may influence neutralization by RBS-C antibodies. Indeed, binding and neutralization of RBS-C antibody CV07-270 was abrogated by RBD-E484K (Figure S1 and Figure 1C). Intriguingly, the five RBS-C antibodies with solved structures are encoded by five different IGHV genes, but target a similar epitope with similar angles of approach. Furthermore, SARS-CoV-2 pseudovirus neutralization by two other highly potent antibodies CC6.29 (*12*) and COVA2-15 (*19*) was markedly reduced by the E484K mutation (Figure 1C). In addition, neutralization by REGN10933 was reduced by both K417N and E484K (Figure 1C), which is a potent antibody used for the therapeutic treatment of COVID-19 (*27*). REGN10933 has a slightly different angle of binding from RBS-A antibodies and tilts slightly towards RBS-B. Thus, K417 can interact with CDRs H1 and H3 of REGN10933, whereas E484 makes contacts with CDR H2 (Figure S11). Overall, our results demonstrate that RBS mutations K417N and E484K can either abolish or extensively reduce the binding and neutralization of several major classes of SARS-CoV-2 RBD antibodies.

**Figure 4.**
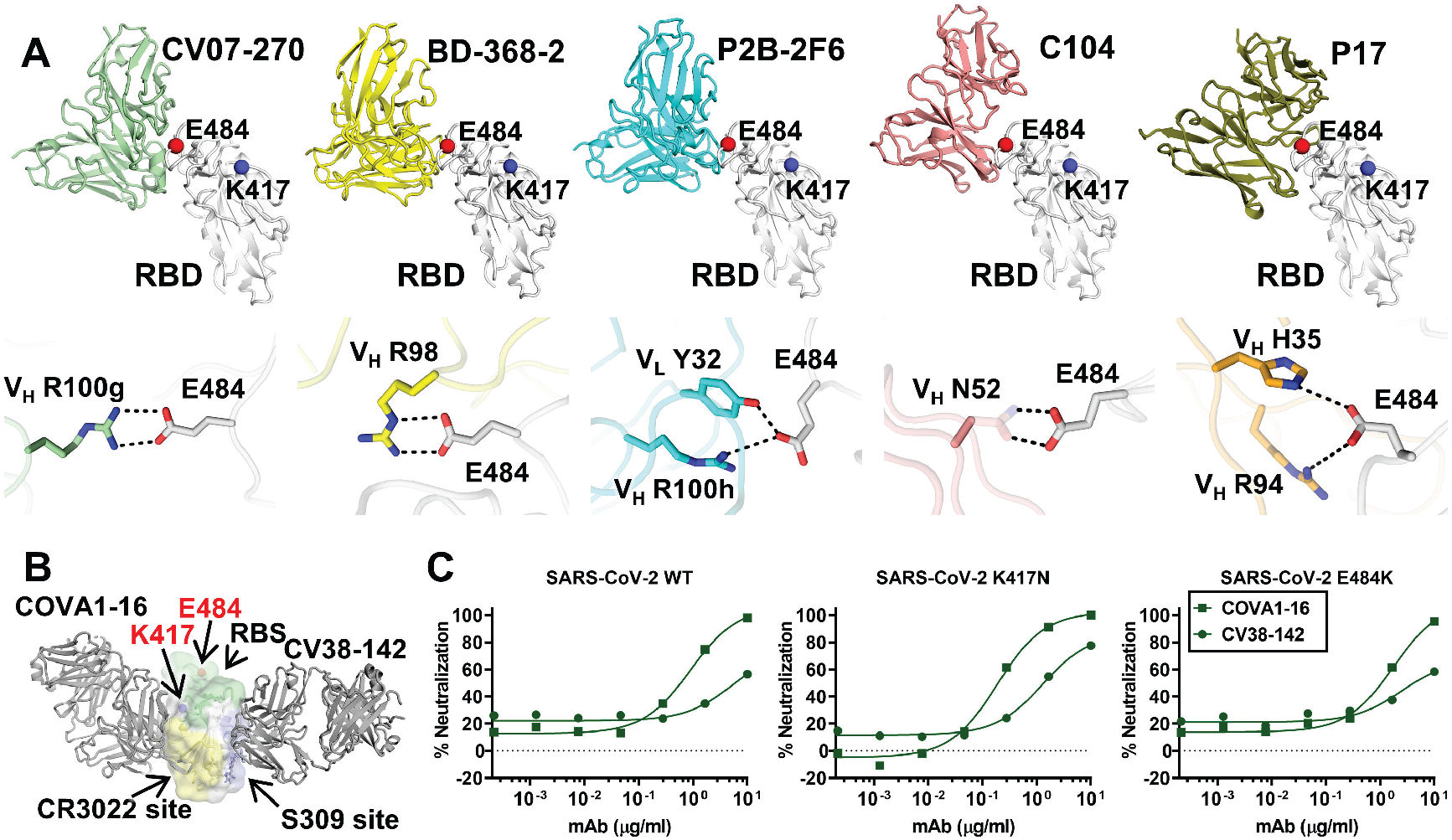
Antibodies targeting other major antigenic sites are differentially affected by mutations in recent variants. **(A)** Interactions between RBS-C antibodies and SARS- CoV-2 RBD. The RBD is shown in white with E484, K417 represented as red and blue spheres, respectively. The various antibodies illustrated are in different colors. Only the variable domains are shown for clarity. Hydrogen bonds and salt bridges to E484 are represented by dashed lines. Published structures with PDB IDs 6XKP (*16*), 7CHF (*37*), 7BWJ (*17*), 7K8U (*44*), and 7CWN (*50*) are used to depict structures of SARS-CoV-2 RBD with CV07-270, BD-368-2, P2B-2F6, C104, and P17, respectively. The electron density for the full side chain of V_H_ N52 was not well resolved in the 3.8-Å structure of C104 in complex with SARS-CoV-2 S. The full side chain is modeled here and shown as transparent sticks to illustrate a possible interaction with E484. **(B)** Cross-neutralizing antibodies to the RBD are not affected by E484 and K417 mutations. COVA1-16 targets the CR3022 cryptic site (yellow) (*51*) and CV38-142 targets the S309 proteoglycan site (blue) (*52*) to the RBD. Glycans at the N343 glycosylation site are represented by sticks. The RBS surface is shown in green. E484 and K417 are highlighted as red and blue spheres, respectively. **(C)** Neutralization of CV38-142 and COVA1-16 against SARS-CoV- 2 wild type, K417N or E484K pseudoviruses.

Two other non-RBS sites that are distant from K417 and E484 have been repeatedly reported to be neutralizing sites on the SARS-CoV-2 RBD, namely the CR3022 cryptic site and S309 proteoglycan site (*46*) (Figures 1C and 4B). We and others have previously shown that antibodies from COVID-19 patients can neutralize SARS-CoV-2 by targeting the CR3022 site, including COVA1-16 (*51*), S304, S2A4 (*30*), and DH1047 (*40*). Recently, we found that CV38-142 also targets the S309 site (*52*). Antibodies targeting these two sites are often cross-reactive with other sarbecoviruses, since these sites are more evolutionarily conserved compared to the RBS. To test the effect of the RBD-K417N and RBD-E484K mutations on neutralizing antibodies that target the S309 and CR3022 sites, we performed binding and neutralization assays on CV38-142 and COVA1-16. Both mutations have minimal effect on these antibodies (Figures 1C and 4C). In fact, recent studies have shown that sera from convalescent or vaccinated individuals can retain neutralization activity, albeit reduced, against the mutated variants (*7, 9, 53*), suggesting that antibodies targeting other epitopes including CR3022 and S309 sites, as well as the NTD, are also present. Thus, the CR3022 cryptic site and S309 site are promising targets to avoid interference by SARS-CoV-2 mutations observed to date.

As SARS-CoV-2 continues to circulate in humans and increasing numbers of COVID-19 vaccines are administered, herd immunity to SARS-CoV-2 is gradually building up both locally and globally. However, as with other RNA viruses, such as influenza and HIV (*54*), further antigenic drift is anticipated in SARS-CoV-2. Such antigenic drift was also observed in at least one well-studied immunosuppressed COVID-19 patient, and included N501Y and E484K mutations (*55*). A recent study also showed the ability of seasonal coronaviruses to undergo antigenic drift (*56*). While antibody responses to the original lineage that initiated COVID-19 pandemic are well characterized in many studies (*12-17, 19, 22, 25, 26, 28*), it is unclear at present how similar the antibody response would be to antigenically distinct lineages. It is possible that IGHV3-53 and IGHV1-2 will not be as enriched in response to infection by the B.1.351 and B.1.1.28.1 lineages or would require greater SHM over the low level reported so far. These questions therefore require urgent attention. Moreover, ongoing efforts to evaluate antibody responses and escape is essential for development and/or modification of COVID-19 vaccines.

In the emerging SARS-CoV-2 lineages, several mutations alter the antigenicity of the RBD as well as the NTD (*57*). While a polyclonal response is generated by natural infection and vaccination, the neutralizing immune response seems to be biased towards particular epitopes and antibody germlines. Many SARS-CoV-2 neutralizing antibodies target the RBS on the RBD, where the most frequent and enriched antibodies are focused on three different sub-epitopes on the RBS. The three most frequent classes (RBS-A, B, C) are adversely affected by mutations at positions 417 and 484 in the South Africa and Brazil lineages. Notwithstanding, some variation in response to these residues from person to person will depend on the characteristics of individual immune responses. However, the epitopes for cross-reactive neutralizing antibodies to the RBD generally do not overlap with the RBS sites. Thus, cross-reactive neutralizing antibodies have not only the potential to confer protection against other zoonotic sarbecoviruses with pandemic potential, but also against antigenically drifted SARS-CoV-2 variants (*58, 59*). As SARS-CoV-2 is likely to become endemic (*60*), it is time to fast track to more broadly effective vaccines and therapeutics that are more resistant to antigenic variation.

## ACKNOWLEDGEMENTS

We thank Henry Tien for technical support with the crystallization robot, Jeanne Matteson and Yuanzi Hua for their contributions to mammalian cell culture, Wenli Yu to insect cell culture, and Robyn Stanfield for assistance in data collection. This work was supported by the Bill and Melinda Gates Foundation OPP1170236 and INV-004923 INV (I.A.W., A.B.W., D.R.B.), NIH R00 AI139445 (N.C.W.), R01 AI132317 (D.N. and D.H.), R01 AI142945 (L.P.), and by the German Research Foundation (H.P.). R.W.S. is a recipient of a Vici fellowship from the Netherlands Organisation for Scientific Research (NWO). This research used resources of the Advanced Photon Source, a U.S. Department of Energy (DOE) Office of Science User Facility, operated for the DOE Office of Science by Argonne National Laboratory under Contract No. DE-AC02-06CH11357. Extraordinary facility operations were supported in part by the DOE Office of Science through the National Virtual Biotechnology Laboratory, a consortium of DOE national laboratories focused on the response to COVID-19, with funding provided by the Coronavirus CARES Act.

## AUTHOR CONTRIBUTIONS

M.Y., D.H., C.C.D.L., N.C.W., and I.A.W. conceived and designed the study. M.Y., C.C.D.L., N.C.W. and H.L. expressed and purified the proteins for crystallization. S.M.R., H.P., and J.K. provided CV05-163 and other antibody clones and sequences. M.J.v.G. and R.W.S, and D.R.B provided plasmids for some of the antibodies reported in (*12, 19*), respectively. M.Y. and X.Z. performed the crystallization, X-ray data collection, determined and refined the X-ray structures. D.H., L.P. and D.N. performed the neutralization assays. A.M.J., and A.B.W. provided nsEM data and performed reconstructions. M.Y., C.C.D.L., N.C.W. and I.A.W. wrote the paper and all authors reviewed and/or edited the paper.

## STRUCTURE DEPOSITIONS

The X-ray coordinates and structure factors have been deposited to the RCSB Protein Data Bank under accession code: 7LOP. The EM maps have been deposited in the Electron Microscopy Data Bank (EMDB) under accession codes: EMD-23466 (one bound), EMD-23467 (two bound), and EMD-23468 (three bound).

## MATERIALS AND METHODS

### Expression and purification of SARS-CoV-2 RBD

Expression and purification of the SARS-CoV-2 spike receptor-binding domain (RBD) were as described previously (*1*). Briefly, the RBD (residues 319-541) of the SARS-CoV- 2 spike (S) protein (GenBank: QHD43416.1) was cloned into a customized pFastBac vector (*2*), and fused with an N-terminal gp67 signal peptide and C-terminal His_6_ tag (*1*). A recombinant bacmid DNA was generated using the Bac-to-Bac system (Life Technologies). Baculovirus was generated by transfecting purified bacmid DNA into Sf9 cells using FuGENE HD (Promega), and subsequently used to infect suspension cultures of High Five cells (Life Technologies) at an MOI of 5 to 10. Infected High Five cells were incubated at 28 °C with shaking at 110 r.p.m. for 72 h for protein expression. The supernatant was then concentrated using a 10 kDa MW cutoff Centramate cassette (Pall Corporation). The RBD protein was purified by Ni-NTA, followed by size exclusion chromatography, and buffer exchanged into 20 mM Tris-HCl pH 7.4 and 150 mM NaCl.

### Expression and purification of Fabs

The heavy and light chains were cloned into phCMV3. The plasmids were transiently co-transfected into ExpiCHO cells at a ratio of 2:1 (HC:LC) using ExpiFectamine™ CHO Reagent (Thermo Fisher Scientific) according to the manufacturer’s instructions. The supernatant was collected at 10 days post-transfection. The Fabs were purified with a CaptureSelect™ CH1-XL Affinity Matrix (Thermo Fisher Scientific) followed by size exclusion chromatography.

### Crystallization and structural determination

A complex of CV05-163 with RBD and CR3022 was formed by mixing each of the protein components at an equimolar ratio and incubating overnight at 4°C. The protein complex was adjusted to 12 mg/ml and screened for crystallization using the 384 conditions of the JCSG Core Suite (Qiagen) on our robotic CrystalMation system (Rigaku) at Scripps Research. Crystallization trials were set-up by the vapor diffusion method in sitting drops containing 0.1 μl of protein and 0.1 μl of reservoir solution. Optimized crystals were then grown in drops containing 0.1 M sodium citrate – citric acid buffer at pH 4.8 and 19% (w/v) polyethylene glycol 6000 at 20°C. Crystals appeared on day 3, were harvested on day 7 by soaking in reservoir solution supplemented with 15% (v/v) ethylene glycol, and then flash cooled and stored in liquid nitrogen until data collection. Diffraction data were collected at cryogenic temperature (100 K) at beamline 23-ID-B of the Advanced Photon Source (APS) at Argonne National Labs with a beam wavelength of 1.033 Å, and processed with HKL2000 (*3*). Structures were solved by molecular replacement using PHASER (*4*). Models for molecular replacement of the RBD and CR3022 were derived from PBD 6W41 (*1*), whereas a model of CV05-163 was generated by Repertoire Builder (https://sysimm.ifrec.osaka-u.ac.jp/rep_builder/) (*5*). Iterative model building and refinement were carried out in COOT (*6*) and PHENIX (*7*), respectively. Epitope and paratope residues, as well as their interactions, were identified by accessing PISA at the European Bioinformatics Institute (http://www.ebi.ac.uk/pdbe/prot_int/pistart.html) (*8*).

### Biolayer interferometry binding assay

Binding assays were performed by biolayer interferometry (BLI) using an Octet Red instrument (FortéBio) as described previously (*1*). To measure the binding kinetics of anti-SARS-CoV-2 IgGs and RBDs (wild type and K417N and E484K variants), the IgGs were diluted with kinetic buffer (1x PBS, pH 7.4, 0.01% BSA and 0.002% Tween 20) into 30 µg/ml. IgG cocktail of two antibodies REGN10933 and REGN10987 were prepared in equimolar ratios of each IgG in 30 µg/ml. The IgGs were then loaded onto anti-human IgG Fc (AHC) biosensors and interacted with 30 µg/ml wild type, K417N, and E484K SARS-CoV-2 RBDs. The assay went through the following steps. 1) baseline: 1 min with 1x kinetic buffer; 2) loading: 150 seconds with IgGs; 3) wash: 1 min wash of unbound IgGs with 1x kinetic buffer; 4) baseline: 1 min with 1x kinetic buffer; 5) association: 2 mins with RBDs; and 6) dissociation: 2 min with 1x kinetic buffer. The binding between IgGs and wild-type RBD acted as a reference for comparison of the binding kinetics to RBD variants.

To obtain kinetics of binding of CV05-163 to SARS-CoV-2 RBD, briefly, His_6_-tagged RBD protein at 20 µg/mL in 1x kinetics buffer (1x PBS, pH 7.4, 0.01% BSA and 0.002% Tween 20) was loaded onto Ni-NTA biosensors and incubated with CV05-163 Fab at concentrations of 500 nM with 2-fold gradient dilutions to 31.25 nM. The assay consisted of five steps: 1) baseline: 60 s with 1x kinetics buffer; 2) loading: 240 s with His_6_-tagged RBD protein; 3) baseline: 60 s with 1x kinetics buffer; 4) association: 180 s with Fab; and 5) dissociation: 180 s with 1x kinetics buffer. For estimating *K*_D_, a 1:1 binding model was used.

### Pseudovirus neutralization assay

Pseudovirus (PSV) preparation and assay were performed as previously described with minor modifications (*9*). Pseudovirions were generated by co-transfection of HEK293T cells with plasmids encoding MLV-gag/pol, MLV-CMV-Luciferase, and SARS-CoV-2 spike WT (GenBank: MN908947) or variants with an 18-AA truncation at the C-terminus. Supernatants containing pseudotyped virus were collected 48 h after transfection and frozen at −80°C for long-term storage. PSV neutralizing assay was carried out as follows. 25 µl of mAbs serially diluted in DMEM with 10% heat-inactivated FBS, 1% Q-max, and 1% P/S were incubated with 25 µl of SARS-CoV-2 PSV at 37°C for 1 h in 96-well half-well plate (Corning, 3688). After incubation, 10,000 Hela-hACE2 cells, generated by lentivirus transduction of wild-type Hela cells and enriched by fluorescence-activated cell sorting (FACS) using biotinylated SARS-CoV-2 RBD conjugated with streptavidin-Alexa Fluor 647 (Thermo, S32357), were added to the mixture with 20 µg/ml Dextran (Sigma, 93556-1G) to enhance infectivity. At 48 h post incubation, the supernatant was aspirated, and HeLa-hACE2 cells were then lysed in luciferase lysis buffer (25 mM Glegly pH 7.8, 15 mM MgSO4, 4 mM EGTA, 1% Triton X-100). Bright-Glo (Promega, PR-E2620) was added to the mixture following the manufacturer’s instruction, and luciferase expression was read using a luminometer. Samples were tested in duplicate, and assays were repeated at least twice for confirmation. Neutralization ID_50_ titers or IC_50_ values were calculated using “One-Site Fit LogIC_50_” regression in GraphPad Prism 9.

### Expression and purification of recombinant S protein for negative-stain electron microscopy

The S protein construct used for negative-stain EM was SARS-CoV-2-6P-Mut7. The construct contains the mammalian-codon-optimized gene encoding residues 1-1208 of the S protein (GenBank: QHD43416.1), an HRV3C cleavage site, and a Twin-strep tag subcloned into the eukaryotic expression vector pcDNA3.4. To prevent cleavage, three amino-acid mutations were introduced into the S1-S2 cleavage site (RRAR to GSAS). For S protein stability, six proline mutations (F817P, A892P, A899P, A942P, K986P, V987P) and a disulfide mutation between T883 and V705 (mutated to cysteines) were introduced (*10–12*). The S plasmid was transfected into HEK293F cells and the supernatant was harvested 6 days post-transfection. To purify the S protein, the supernatant was run through a Stereotactic XT 4FLOW column (IBA Lifesciences) followed by size exclusion chromatography using a Superose 6 increase 16/600 pg. column (GE Healthcare Biosciences). Protein fractions corresponding to the trimeric S protein were collected and concentrated.

### nsEM sample preparation and data collection

SARS-CoV-2-6P-Mut7 S protein was complexed with a 3-fold molar excess of Fab and incubated for 30 minutes at room temperature. The complex was diluted to approximately 0.03 mg/mL with 1x TBS pH 7.4 and applied onto carbon-coated 400- mesh copper grids. The grids were stained with 2% (w/v) uranyl-formate for 60 seconds immediately following sample application. Grids were imaged at 200 keV on a FEI Tecnai T20 using a Tietz TVIPS CMOS 4k × 4k camera at 62,000× magnification, −1.50 μm defocus, and a total dose of 25 e^−^/Å^2^. Micrographs were collected using Leginon (*13*) and transferred to the Appion database (*14*) for processing. Particles were picked using a difference-of-Gaussians picker (DoG-picker) (*15*) and stacked with a box size of 256 pixels, then transferred to Relion (*16*) for 2D and 3D classification. Select 3D classes were refined and analyzed in UCSF Chimera (*17*) for making figures. A published prefusion spike model (PDB: 6VYB) (*18*) was used in the structural analysis.

---

**Supplementary Figure 1.**
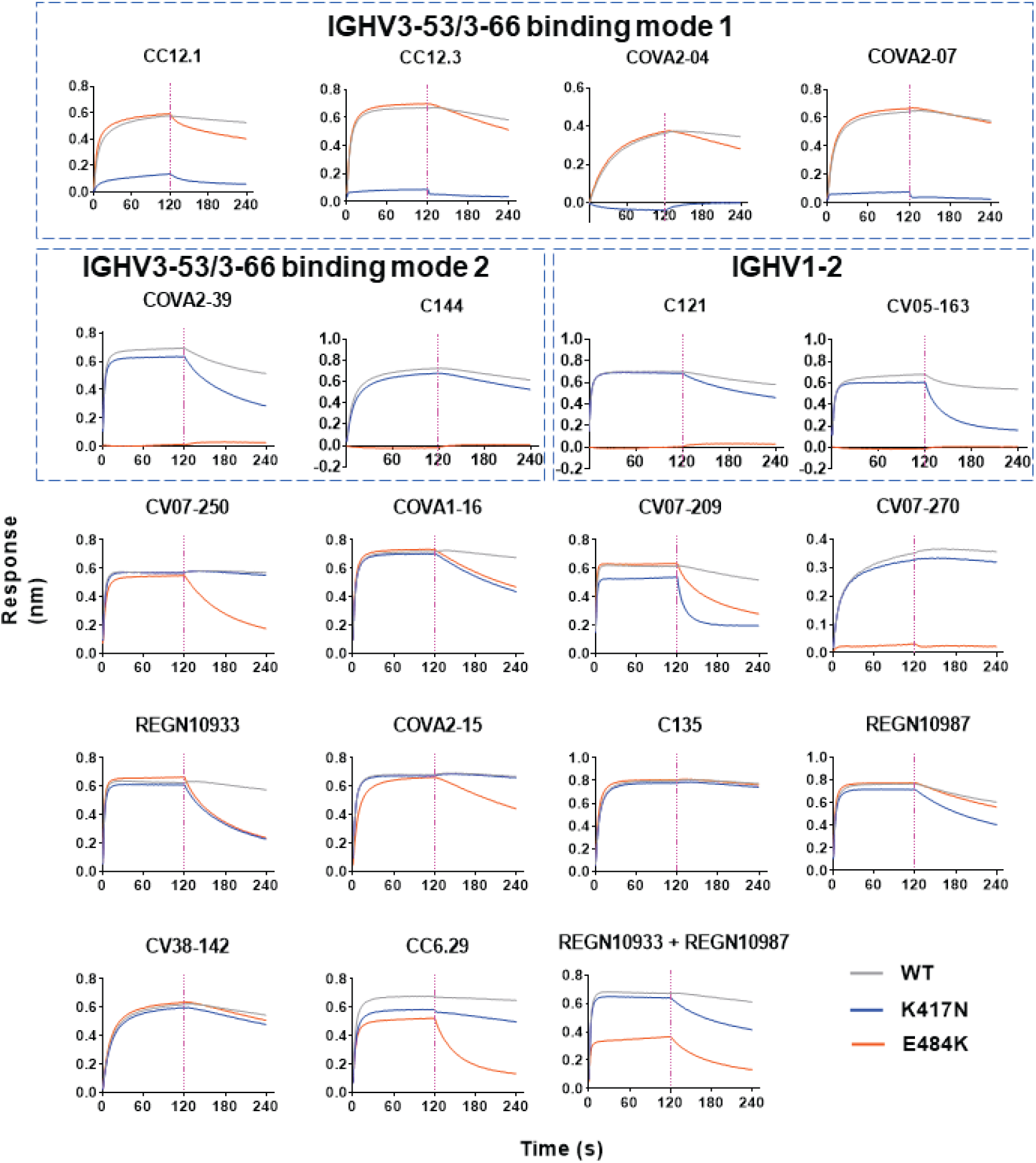
Sensorgrams for binding of IgGs to wild-type and mutated SARS-CoV-2 RBD. Binding kinetics were measured by biolayer interferometry with IgGs loaded on the biosensor and RBD proteins in solution. Wild-type SARS-CoV-2 RBD, K417N, and E484K are shown in grey, blue, and orange, respectively. Representative results of three replicates for each experiment are shown.

**Supplementary Figure 2.**
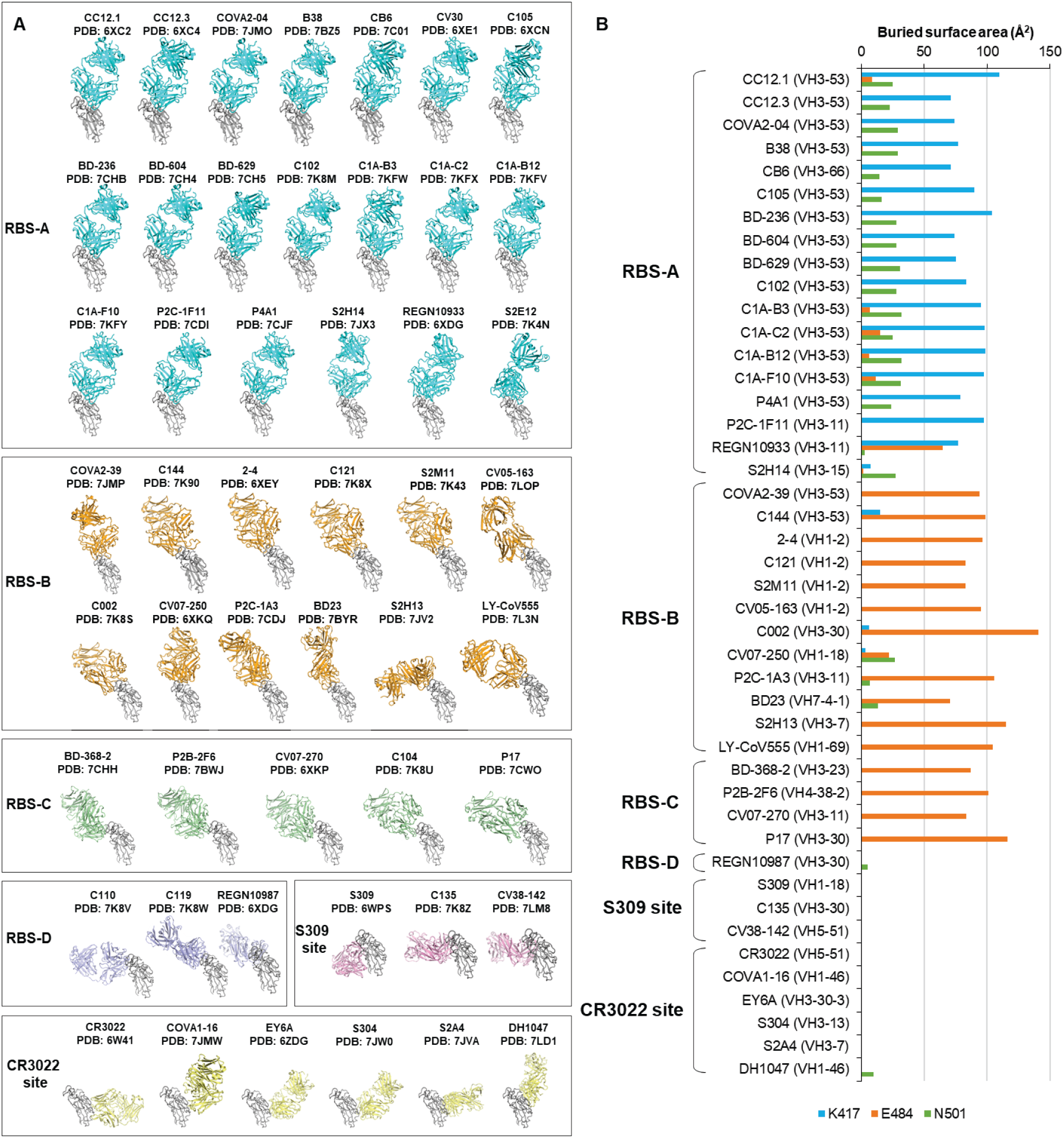
Epitope classification of RBD-targeting antibodies. **(A)** The classification is based on initial categories and assignments in (*19*). The SARS-CoV-2 RBD (grey) is shown in the same relative orientation in each panel. Antibodies are color-coded by their respective epitopes. **(B)** Buried surface area (BSA) of K417, E484, and N501 of the SARS-CoV-2 RBD by SARS-CoV-2 targeting antibodies as calculated by the PISA program (*8*). PDB codes for the structures used for the BSA calculation are shown in panel (A). Structures of S2E12 (PDB 7K4N), C104 (PDB 7K8U), C110 (PDB 7K8V), and C119 (PDB 7K8W) are not included in the calculation because residues or side chains of the epitope and/or paratope residues were truncated in the published structures. Heavy chain germline genes that encode each antibody are shown in brackets.

**Supplementary Figure 3.**
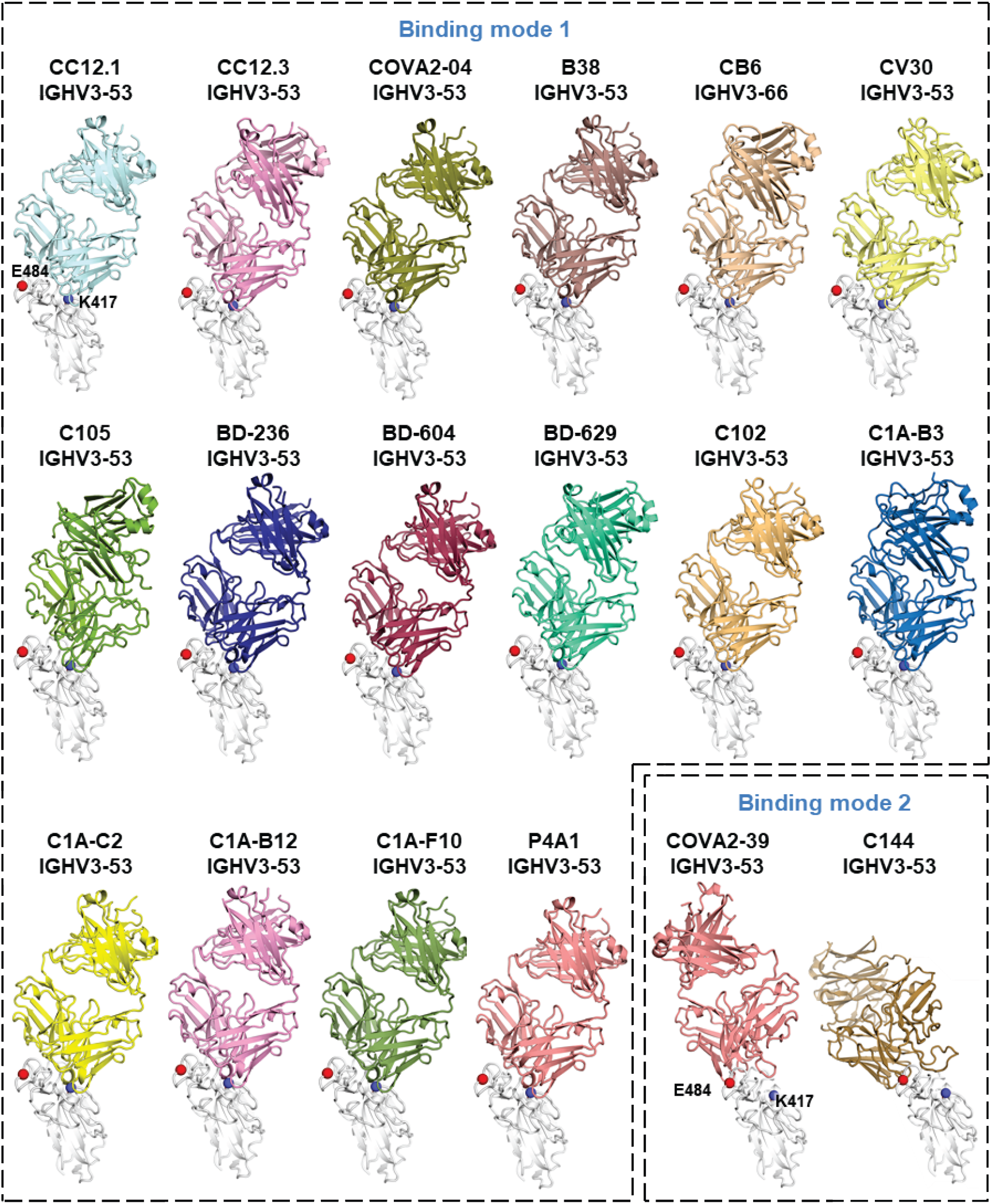
Structures of IGHV3-53/3-66 antibodies in complex with SARS-CoV-2 RBD. 16 out of 18 RBD-targeting IGHV3-53/3-66 antibodies with available structures in the PDB bind to the same epitope using a nearly identical angle of approach (binding mode 1) to SARS-CoV-2 RBD (white). RBD-K417 is intimately involved in the epitope. The other two antibodies COVA2-39 and C144 bind to the opposite site of the RBS (binding mode 2) and, in this binding mode, RBD-E484 is a key contributor to the epitope. RBDs are shown in the same relative orientation in each panel. E484 (left) and K417 (right) are represented by red and blue spheres, respectively, and are also labeled in the first panel. All available RBD-targeting IGHV3- 53/3-66 antibody structures in the PDB at time of analysis (January 2021) are shown: CC12.1 (PDB ID: 6XC3), CC12.3 (PDB ID: 6XC4) (*20*), COVA2-04 (PDB ID: 7JMO) (*21*), B38 (PDB ID: 7BZ5) (*22*), CB6 (PDB ID: 7C01) (*23*), CV30 (PDB ID: 6XE1) (*24*), C105 (PDB ID: 6XCN) (*25*), BD-236 (PDB ID: 7CHB), BD-604 (PDB ID: 7CH4), BD-629 (PDB ID: 7CH5) (*26*), C102 (PDB ID: 7K8M) (*27*), C1A-B3 (PDB ID: 7KFW), C1A-C2 (PDB ID: 7KFX), C1A-B12 (PDB ID: 7KFV), C1A-F10 (PDB ID: 7KFY) (*28*), P4A1 (PDB ID: 7JCF) (*29*), COVA2-39 (PDB ID: 7JMP) (*21*), and C144 (PDB ID: 7K90) (*25*).

**Supplementary Figure S4.**
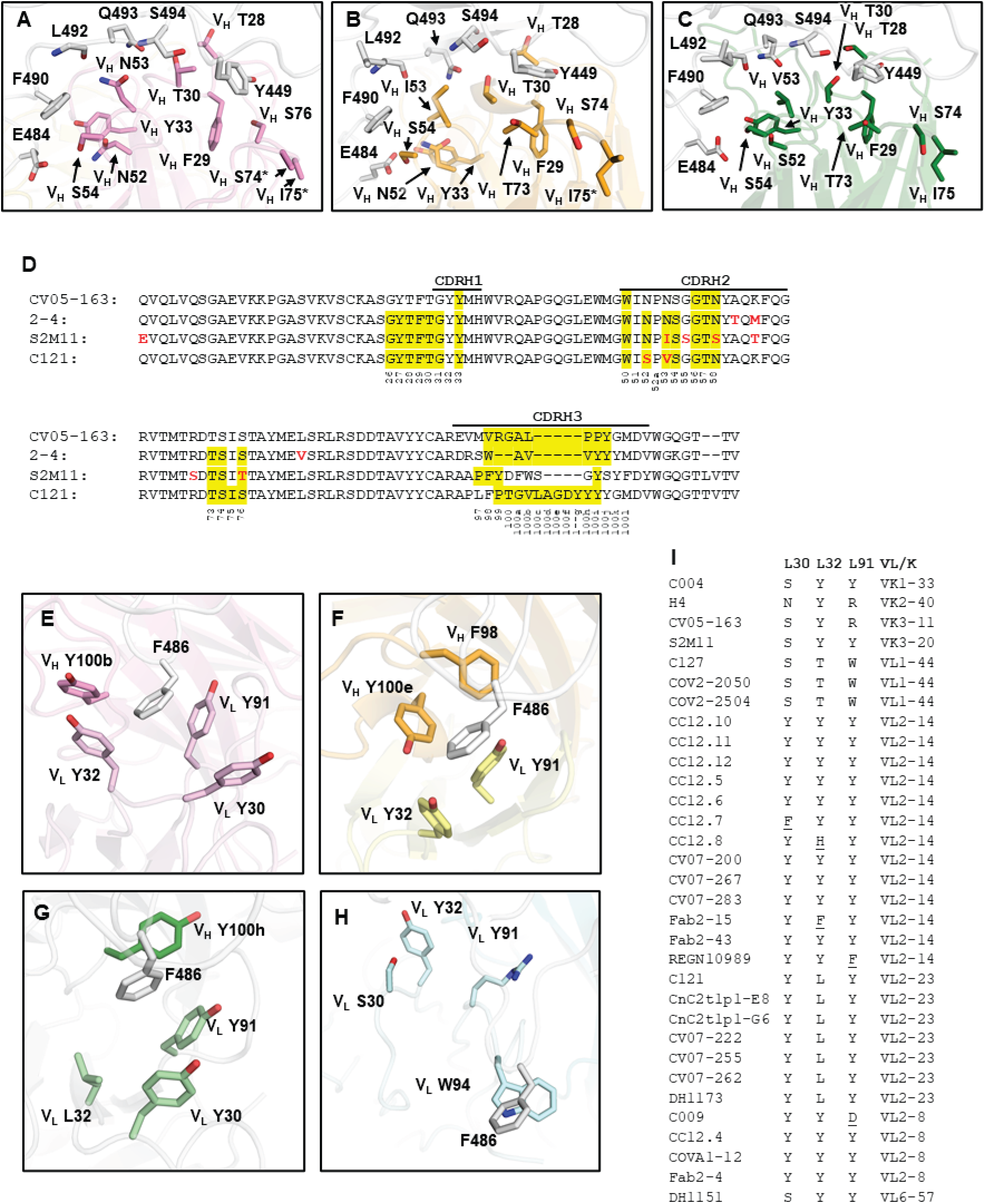
Molecular interactions between IGHV1-2 antibodies and SARS-CoV-2 RBD. In panels A-C and E-H, RBD residues are shown in white. Details of antibody interactions with the RBS of the SARS-CoV-2 RBD are illustrated for: **(A)** 2-4 (PDB 6XEY) (*30*) in pink, **(B)** S2M11 (PDB 7K43) (*31*) in orange, and **(C)** C121 (PDB 7K8X) (*27*) in green. **(D)** Sequence alignment of CV05-163, 2-4, and S2M11 variable heavy (V_H_ region). The regions that correspond to CDR H1, H2, H3, L1, L2, and L3 are indicated in Kabat numbering. Antibody residues that interact with the RBD are highlighted in yellow [residues with a BSA > 0 Å^2^ as calculated by the PISA program (*8*)]. Somatic hypermutated residues are highlighted in red. **(E–H)** Interactions of SARS-CoV- 2 RBD F486 with **(E)** 2-4, **(F)** S2M11, **(G)** C121, and **(H)** CV05-163. The four structures are superimposed on the RBD and shown in the same overall view. **(I)** Light-chain residues at positions 30, 32, and 91 (Kabat numbering) in all RBD-targeting IGHV1-2 neutralizing antibodies with sequence information [summarized in CoV-AbDab (*32*)]. Somatically hypermutated residues are underlined. Among IGHV1-2 RBD antibodies with reported neutralization activity, nine different light chains have been observed to date, although with preference for IGLV2-14, IGLV2-23, and IGLV2-8, which account for over 76% of the light chains that pair with IGHV1-2. Importantly, germline residues at positions 30, 32, and 91 of IGLV2-14, IGLV2-23, and IGLV2-8 are all aromatic or hydrophobic residues, further delineating why IGHV1-2 antibodies paired with these specific light chains are naturally favored for this canonical RBD-binding mode for neutralization of SARS-CoV-2. In contrast, CV05-163 represents a small subset of IGHV1-2 RBD-targeting antibodies without hydrophobic residues at positions 30 or 91 of the light chain that form a hydrophobic pocket for anchoring RBD-F486 in most IGHV1-2 antibodies that bind in the canonical mode.

**Supplementary Figure 5.**
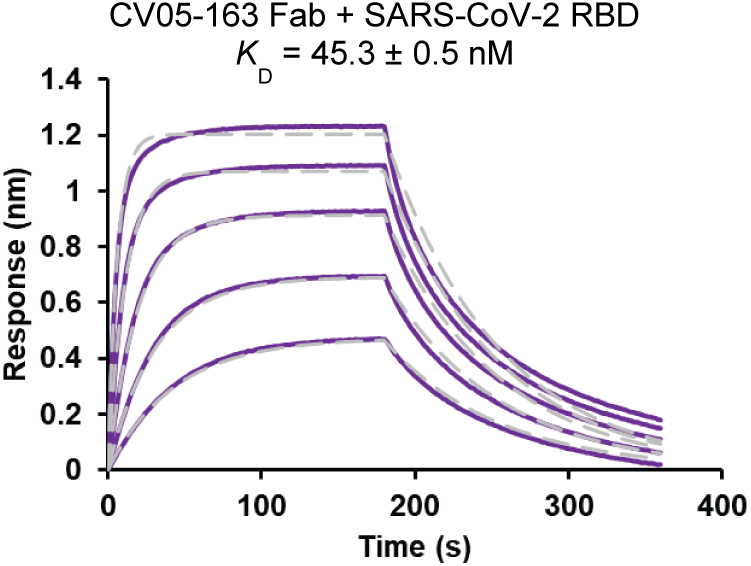
Sensorgrams for binding of CV05-163 Fab to SARS-CoV- 2 RBD. Binding kinetics of CV05-163 Fab against SARS-CoV-2 RBD were measured by biolayer interferometry (BLI). Y-axis represents the response. Purple solid lines represent the response curves and grey dashed lines represent the 1:1 binding model. Binding kinetics were measured for five concentrations of Fab at 2-fold dilution ranging from 500 nM to 31.25 nM. The K_D_ of the fitting is indicated. Representative results of three replicates for each experiment are shown.

**Supplementary Figure 6.**
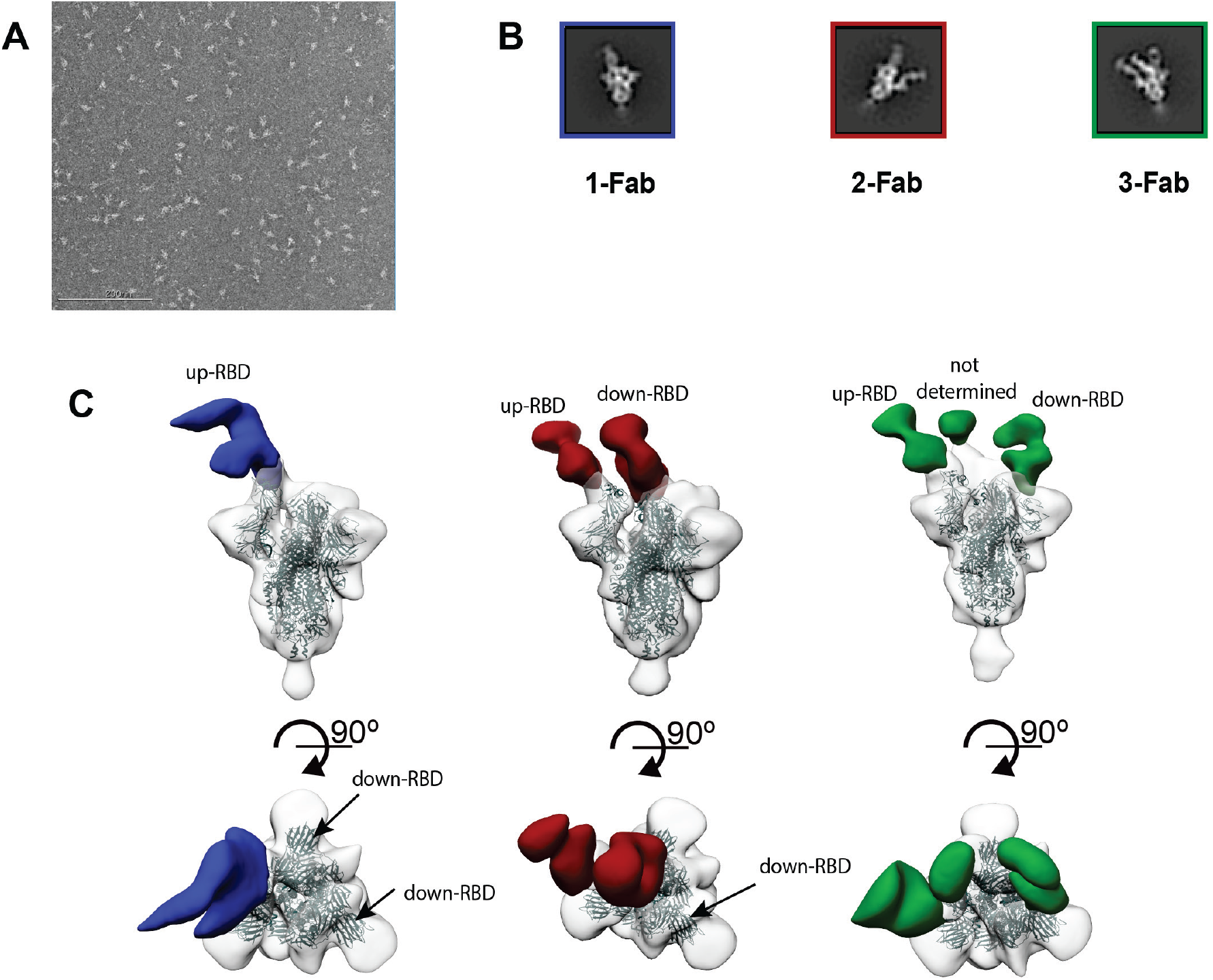
nsEM analysis of CV05-163 in complex with SARS-CoV-2 S trimer. **(A)** Representative negative stain-EM micrograph. **(B)** Select 2D class averages of single-particle nsEM analysis of CV05-163 complexed with S trimer. 2D classes corresponding to the 1-, 2-, and 3-Fab binding stoichiometries are highlighted in a blue, red, and green box, respectively. **(C)** 3D nsEM reconstructions of 1, 2, and 3 Fab CV05- 163 bound to the SARS-CoV-2 S trimer. CV05-163 binds both up- and down-RBD in various stoichiometries, including molar ratios of 1:1, 2:1, and 3:1 (Fab : S trimer).

**Supplementary Figure 7.**
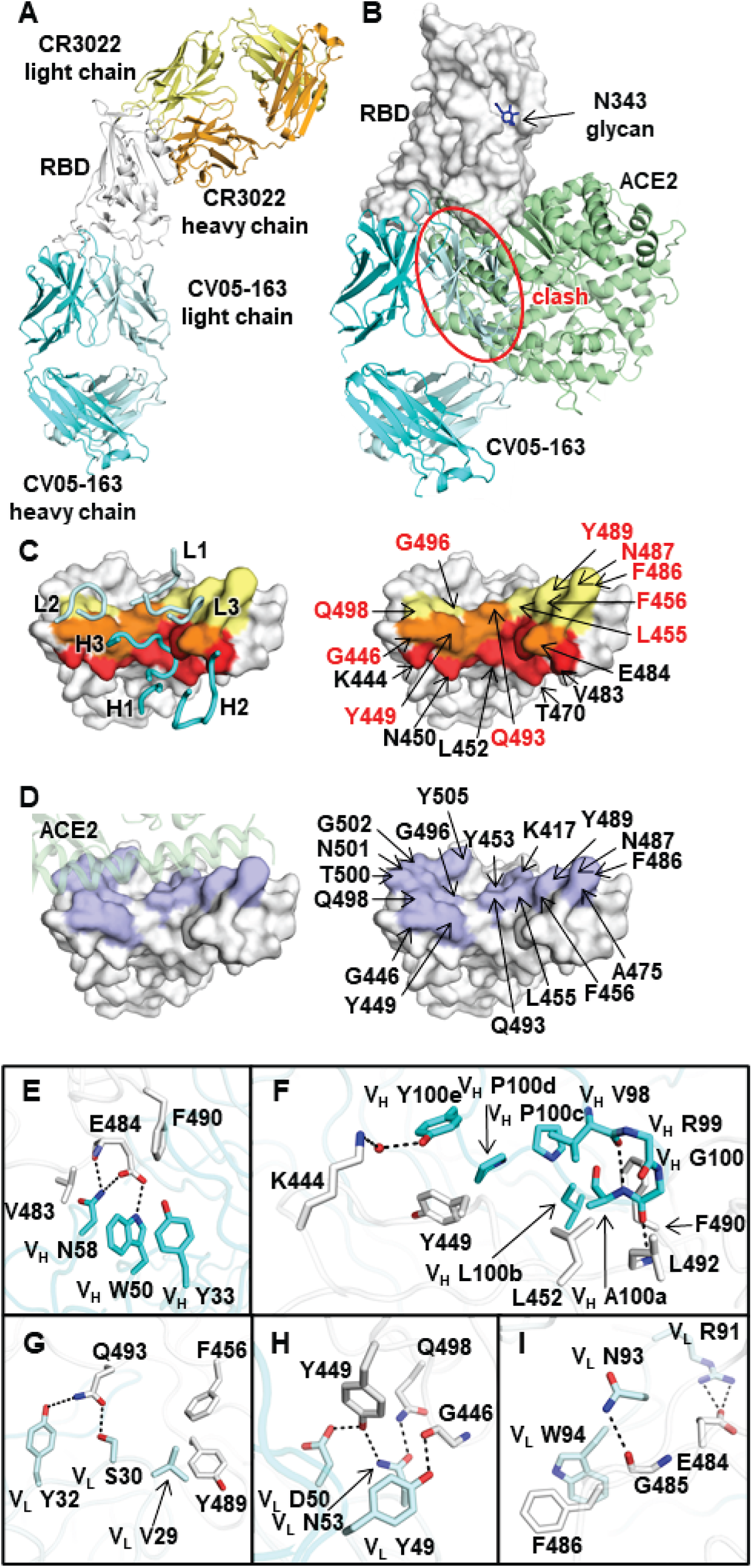
Crystal structure of SARS-CoV-2 RBD in complex with Fabs CV05-163 and CR3022. **(A)** The binding site of CV05-163 (Fab heavy and light chains shown in cyan and pale cyan, respectively) on the RBD (white) is distinct from that of CR3022 (Fab heavy and light chains shown in orange and yellow, respectively). **(B)** The ACE2/RBD complex structure (PDB 6M0J) (*33*) is superimposed on the CV05-163/RBD complex. CV05-163 (cyan) would clash with ACE2 (green) if bound simultaneously with the RBD (indicated by red ellipse). The N-glycan at N343 of the RBD is in dark blue. **(C)** Epitope of CV05-163. Epitope residues contacting the heavy chain are in red and the light chain in yellow, while residues contacting both heavy and light chains are in orange. On the left panel, CDR loops are labeled. On the right panels, epitope residues are labeled. For clarity, only representative epitope residues are labeled. Epitope residues that are also involved in ACE2 binding are labeled in red. **(D)** ACE2-binding residues on the RBD are in lilac. On the left panel, ACE2 is represented by an olive semi-transparent surface. On the right panel, ACE2-binding residues are labeled. The 17 ACE2-binding residues are as described previously (PDB 6M0J) (*33*). **(E and F)** Interactions between the RBD and heavy chain of CV05-163 for **(E)** CDR H1 and H2 and **(F)** CDR H3. **(G to I)** Interactions between RBD and light chain of CV05-163 for **(G)** CDR L1, **(H)** CDR L2 and **(I)** CDR L3.

**Supplementary Figure 8.**
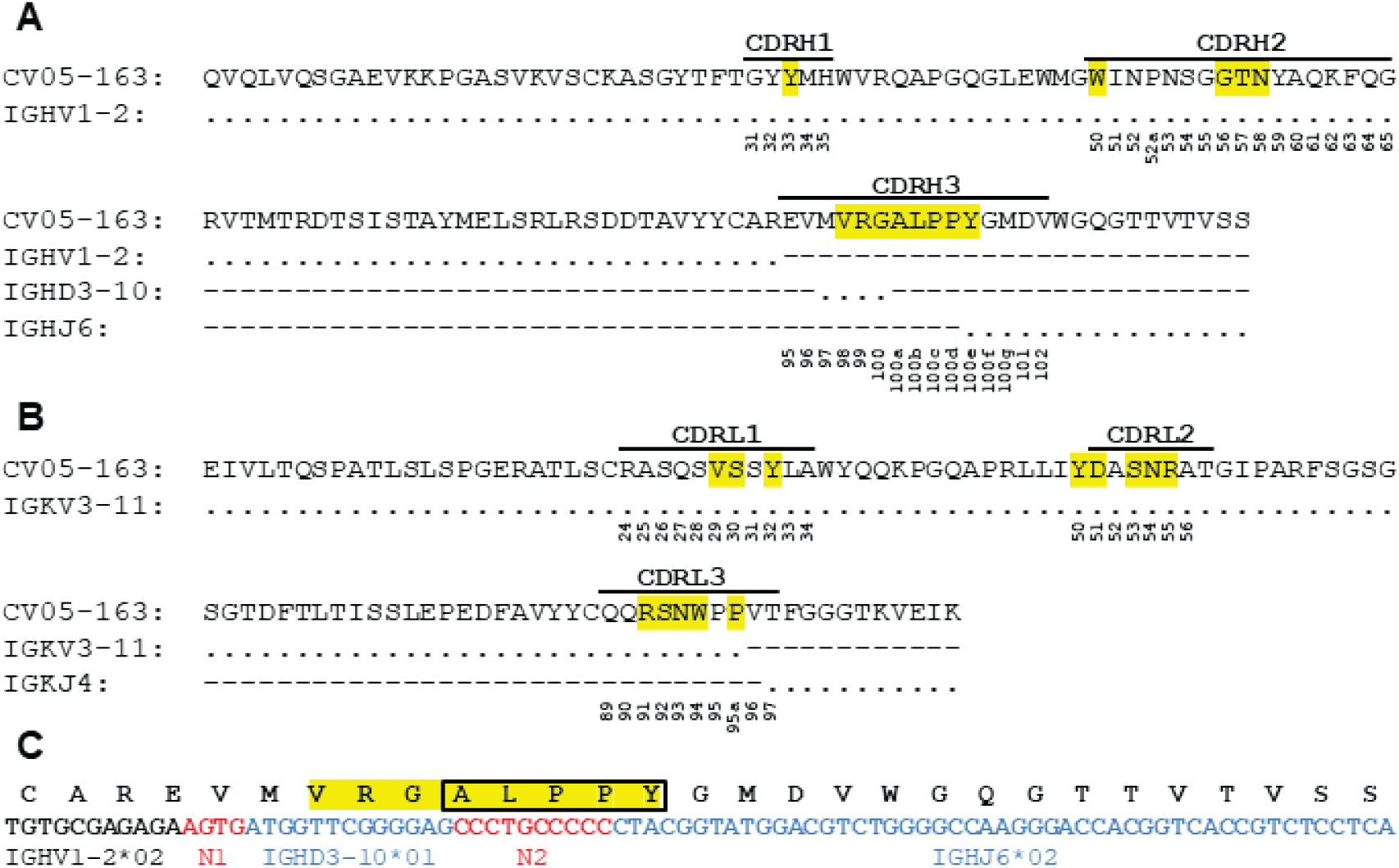
Sequence analysis of a germline antibody CV05-163 targeting the SARS-CoV-2 RBD. (A and B) CV05-163 V_H_ and V_L_ sequence alignment with corresponding putative germline gene segments. The sequences of both V_H_ and V_L_ chains of CV05-163 have no somatic hypermutations compared to their germline genes. The regions that correspond to CDR H1, H2, H3, L1, L2, and L3 are indicated. Residues that interact with the RBD are highlighted in yellow. Residue positions in the CDRs are labeled according to the Kabat numbering scheme. Conserved residues are represented by dots (.) and non-germline encoded residues are indicated by dashes (-). **(C)** Sequence of the V-D-J junction of CV05-163, with putative gene segments (blue) and N-regions (red) indicated. Residues that interact with the RBD are highlighted in yellow. The shared junctional motif in CDR H3 (“ALPPY”) arises mainly from N-additions and is highlighted in a black box.

**Supplementary Figure 9.**
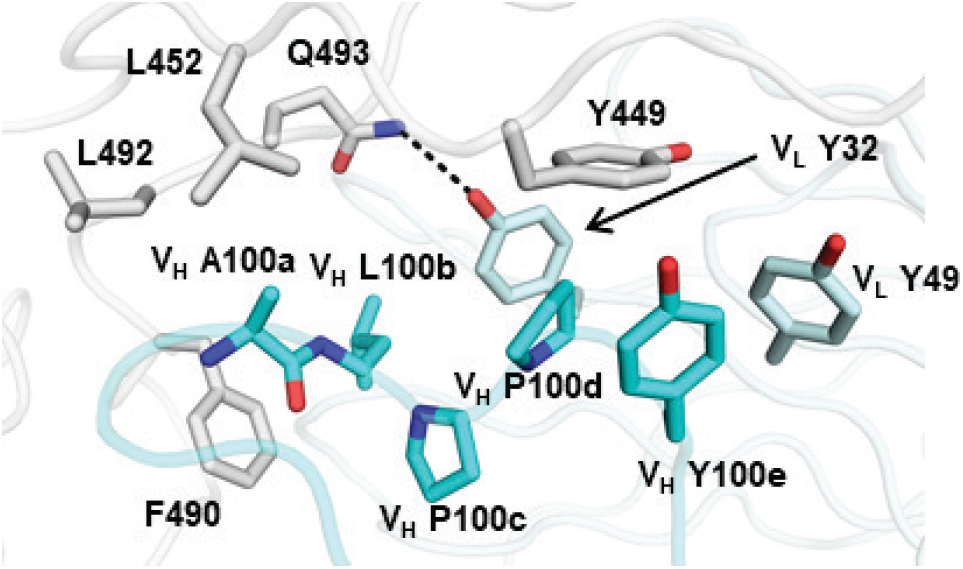
Interactions between the ALPPY motif and SARS-CoV-2 RBD. The RBD is in white and antibody residues in cyan (heavy chain) and pale cyan (light chain), respectively. A hydrogen bond is represented by a dashed line.

**Supplementary Figure 10.**
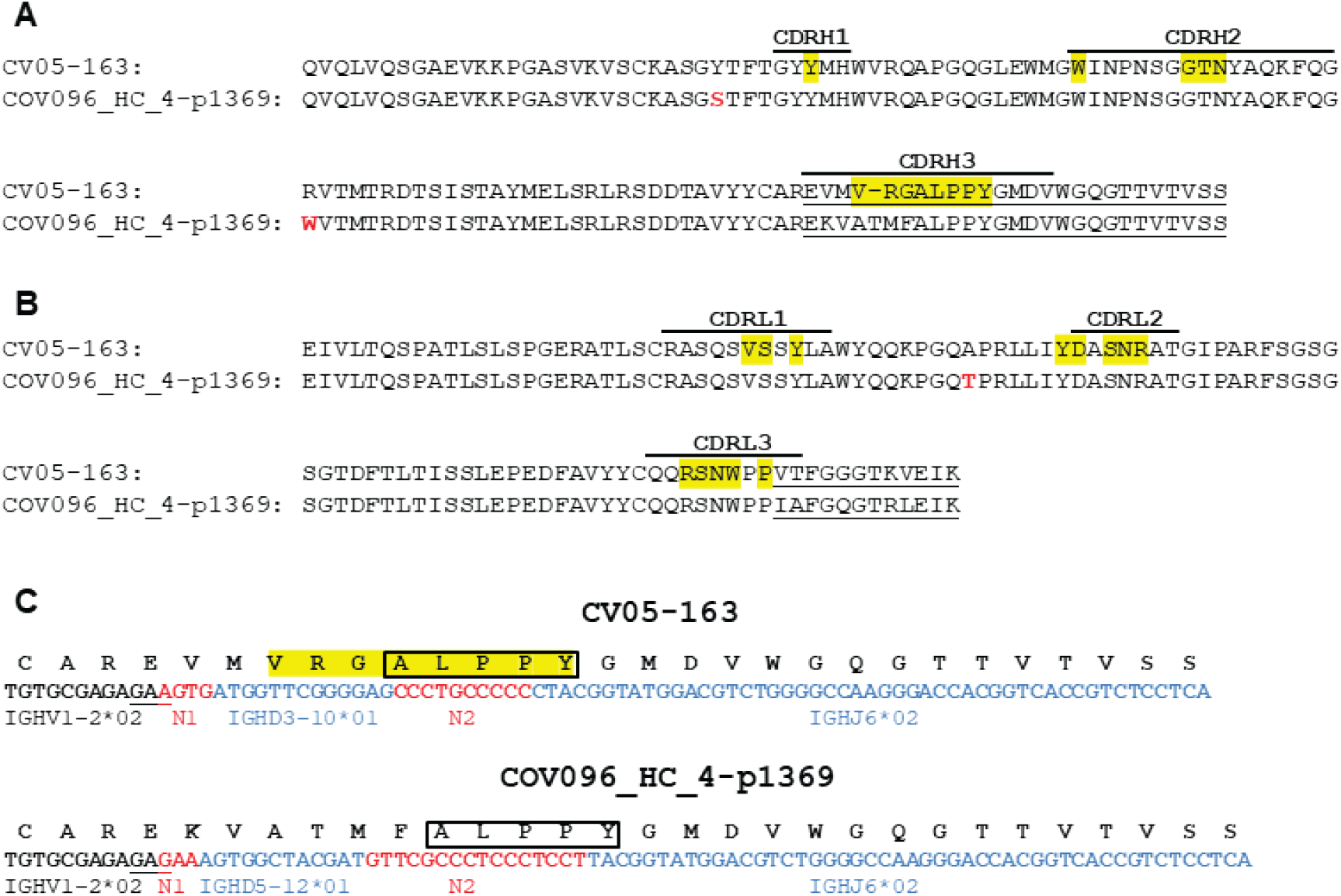
Convergence of the N-addition encoded ALPPY motif in RBD-targeting IGHV1-2 antibodies. Conservation of a non-germline encoded motif is expected to be extremely rare, but an “ALPPY” motif encoded mainly by N-additions is found in CDR H3 of two RBD antibodies CV05-163 and COV096_HC_4-p1369 (*34*) from different individuals. CV05-163 and COV096_HC_4-p1369 are both encoded by IGHV1- 2/IGKV3-11. **(A and B)** V_H_ and V_L_ sequence alignment of CV05-163 with COV096_HC_4-p1369. Somatic hypermutated residues are highlighted in red. Regions that correspond to CDR H1, H2, H3, L1, L2, and L3 are indicated. Residues of CV05- 163 that interact with the RBD in the CV05-163/RBD complex structure are highlighted in yellow [defined here as residues with a BSA > 0 Å^2^ as calculated by the PISA program (*8*)]. Residue positions in the CDRs are labeled according to the Kabat numbering scheme. Non-V-gene encoded residues are underlined. **(C)** Sequences of the V-D-J junction of CV05-163 and COV096_HC_4-p1369, with putative D and J gene segments (blue) and N-regions (red) indicated. CV05-163 residues that interact with the RBD are highlighted in yellow. The shared junctional motif (“ALPPY”) is boxed. Germline residue V_L_ R91 of CV05-163 that interacts with RBD-E484 is also stabilized by V_H_ E95, which is also partially encoded by N-additions and conserved between CV05-163 and COV096_HC_4-p1369. Nucleotides that encode V_H_ E95 of CV05-163 and COV096_HC_4-p1369 are underlined.

**Supplementary Figure 11.**
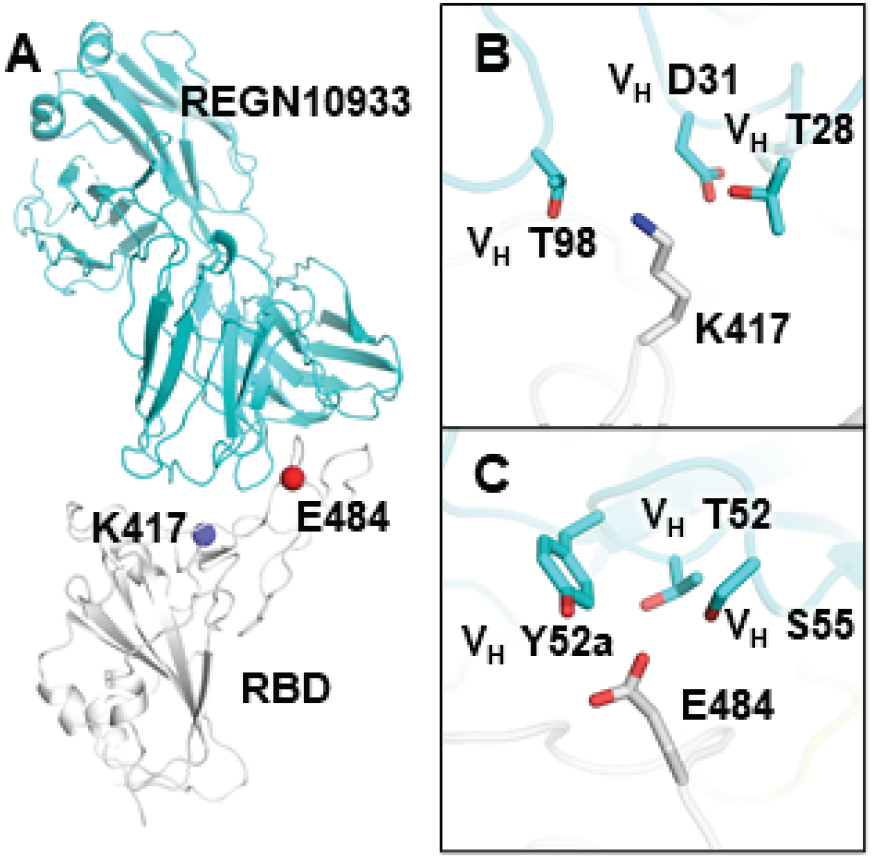
K417 and E484 are both involved in recognition by antibody REGN10933. **(A)** An overall view of the interaction between REGN10933 (cyan) and SARS-CoV-2 RBD (white) (PDB 6XDG) (*35*). Residues K417 and E484 are represented by blue and red spheres, respectively. **(B-C)** Interactions of **(B)** RBD-K417 and **(C)** RBD-E484 with REGN10933. Hydrogen bonds are not shown due to the limited resolution (3.9 Å). Kabat numbering is assigned to the residues of REGN10933.

**Supplementary Table 1.** X-ray data collection and refinement statistics.

|  |  |
| --- | --- |
| **Data collection** | CV05-163 + RBD + CR3022 |
| Beamline | APS23ID-B |
| Wavelength (Å) | 1.0337 |
| Space group | P 1 2<sub>1</sub> 1 |
| Unit cell parameters |  |
| a, b, c (Å) | 86.0, 132.9, 111.3 |
| α, β, γ (°) | 90, 100.7, 90 |
| Resolution (Å)<sup>a</sup> | 50.0-2.25 (2.29-2.25) |
| Unique reflections<sup>a</sup> | 115,419 (10,628) |
| Redundancy<sup>a</sup> | 3.2 (2.4) |
| Completeness (%)<sup>a</sup> | 98.9 (91.5) |
| <I/σ<sub>I</sub>><sup>a</sup> | 7.1 (1.0) |
| R<sub>sym</sub><sup>b</sup> (%)<sup>a</sup> | 14.6 (63.8) |
| R<sub>pim</sub><sup>b</sup> (%)<sup>a</sup> | 9.4 (46.6) |
| CC<sub>1/2</sub><sup>c</sup> (%)<sup>a</sup> | 97.8 (61.5) |
| **Refinement statistics** |  |
| Resolution (Å) | 49.4-2.25 |
| Reflections (work) | 115,390 |
| Reflections (test) | 10,625 |
| R<sub>cryst</sub><sup>d</sup> / R<sub>free</sub><sup>e</sup> (%) | 22.1/26.5 |
| No. of atoms | 17,694 |
| RBD | 3,101 |
| Fab | 13,328 |
| Glycan | 28 |
| Solvent | 1,237 |
| Average B-values (Å<sup>2</sup>) | 34 |
| RBD | 31 |
| CV05-163 Fab | 33 |
| CR3022 Fab | 35 |
| Glycan | 47 |
| Solvent | 35 |
| Wilson B-value (Å<sup>2</sup>) | 29 |
| **RMSD from ideal geometry** |  |
| Bond length (Å) | 0.004 |
| Bond angle (°) | 0.82 |
| **Ramachandran statistics (%)** |  |
| Favored | 97.1 |
| Outliers | 0.09 |
| **PDB code** | 7LOP |

<sup>a</sup> Numbers in parentheses refer to the highest resolution shell.
<sup>b</sup> $R_{sym} = \sum_{hkl} \sum_i |I_{hkl,i} - \langle I_{hkl} \rangle| / \sum_{hkl} \sum_i I_{hkl,i}$ and $R_{pim} = \sum_{hkl} (1/(n-1))^{1/2} \sum_i |I_{hkl,i} - \langle I_{hkl} \rangle| / \sum_{hkl} \sum_i I_{hkl,i}$ , where $I_{hkl,i}$ is the scaled intensity of the $i^{th}$ measurement of reflection h, k, l, $\langle I_{hkl} \rangle$ is the average intensity for that reflection, and n is the redundancy.
<sup>c</sup> CC<sub>1/2</sub> = Pearson correlation coefficient between two random half datasets.
<sup>d</sup> $R_{cryst} = \sum_{hkl} |F_o - F_c| / \sum_{hkl} |F_o| \times 100$ , where $F_o$ and $F_c$ are the observed and calculated structure factors, respectively.
<sup>e</sup> $R_{free}$ was calculated as for $R_{cryst}$ , but on a test set comprising 5% of the data excluded from refinement.

**Supplementary Table 2.** Hydrogen bonds and salt bridges identified at the antibody-RBD interface using the PISA program.

| SARS-CoV-2<br>RBD | Distance<br>[Å] | CV05-163 |
| --- | --- | --- |
| **Hydrogen bonds** |  |  |
| GLU484[OE2] | 3.0 | VH TRP50[NE1] |
| GLU484[O] | 3.0 | VH ASN58[ND2] |
| LEU492[N] | 2.7 | VH GLY100[O] |
| GLY446[O] | 2.7 | VK: TYR49[OH] |
| TYR449[OH] | 3.4 | VK: ASN53[ND2] |
| GLU484[OE2] | 3.0 | VK: ARG91[NH1] |
| GLU484[OE2] | 3.1 | VK: ARG91[NH2] |
| GLY485[O] | 3.3 | VK: ASN93[ND2] |
| GLN493[OE1] | 2.9 | VK: SER30[OG] |
| GLN493[NE2] | 3.7 | VK: TYR32[OH] |
| TYR449[OH] | 2.5 | VK: ASP50[OD2] |
| GLN498[NE2] | 2.7 | VK: ASN53[OD1] |
| **Salt bridges** |  |  |
| GLU484[OE2] | 3.0 | VK: ARG91[NH1] |
| GLU484[OE2] | 3.1 | VK: ARG91[NH2] |

